# Emergence of linkage between cooperative RNA replicators encoding replication and metabolic enzymes thorough experimental evolution

**DOI:** 10.1101/2022.10.11.511852

**Authors:** Kensuke Ueda, Ryo Mizuuchi, Norikazu Ichihashi

## Abstract

The integration of individually replicating genes into a primitive chromosome is a key evolutionary transition in the development of life, allowing the simultaneous inheritance of genes. However, how this transition occurred is unclear because of the extended size of primitive chromosomes, which replicate slower than unlinked genes. Theoretical studies have suggested that a primitive chromosome can evolve in the presence of cell-like compartments, as the physical linkage prevents the stochastic loss of essential genes upon division, but experimental support for this is lacking. Here, we demonstrate the evolution of a chromosome-like RNA from two cooperative RNA replicators encoding replication and metabolic enzymes. Through their long-term replication in cell-like compartments, linked RNAs emerged with the two cooperative RNAs connected end-to-end. The linked RNAs had different mutation patterns than the two unlinked RNAs, suggesting that they were maintained as partially distinct lineages in the population. Our results provide experimental evidence supporting the plausibility of the evolution of a primitive chromosome from unlinked gene fragments, an important step in the emergence of complex biological systems.

**Author Summary:** The integration of genes into a chromosome is a fundamental genetic organization in all extant life. The assembly of unlinked genes during prebiotic evolution was likely a major evolutionary transition toward the development of a complex cell. Decades of theoretical studies have suggested a plausible evolutionary pathway to a primitive chromosome from replicating RNA molecules that harbor cooperative genes within a protocell structure. However, demonstrating the evolution of a primitive chromosome in an experimental setup is challenging. We previously developed a cooperative RNA replication system in which two types of RNAs co-replicate using their self-encoded replication and metabolic enzymes. Using this system, in the present study, we demonstrate the evolution of a linkage between the two cooperative RNA replicators in compartments. An evolved “linked” RNA harbored the entire region of both genes, accumulated distinct mutations, and retained the ability to replicate using the two proteins translated from itself. These experimental findings support a prebiotic evolutionary scenario, in which unlinked genes assembled into a single genomic structure.

## Introduction

All extant cells have chromosomes that harbor multiple genes and ensure their organized inheritance. In the early evolution of life, such a genome organization may have been absent; unlinked RNA molecules encoding different functions or genes cooperated with each other for replicating the entire system [1–6]. The subsequent appearance of a linkage between cooperative RNA replicators, or a primitive chromosome, is considered a major evolutionary transition toward complex biological organization [2–4]. Chromosome formation could have been both advantageous and disadvantageous for primitive life. Chromosomes enabled the synchronized replication of essential genes and potentially drove the evolution of efficient enzymes [7]. However, chromosomes were longer than unlinked RNA and hence required more time to replicate and had a higher chance of degradation. It remains unclear whether a primitive chromosome could evolve from individual cooperating RNA molecules despite these disadvantages.

Theoretical studies have suggested that the presence of primitive cell-like compartments (protocells) could have facilitated the selection of a primitive chromosome over individual RNA replicators [8–11]. The random assortment of cooperatively replicating RNAs upon protocell division may have caused the stochastic loss of cooperating partners. In contrast, the formation of a primitive chromosome ensured the co-encapsulation of cooperative genes, and therefore, a chromosome could have had an evolutionary advantage over unlinked replicators in compartments [8,9,11]. Although previous studies have established theoretical frameworks for the evolution of a primitive chromosome, empirical demonstration is lacking.

Previously, we constructed an experimental cooperative RNA replication system consisting of two RNAs and a reconstituted translation system [12]. One of the RNAs encodes a subunit of Qβ replicase, an RNA-dependent RNA polymerase derived from Qβ phage, and the other encodes a metabolic enzyme, nucleotide diphosphate kinase (NDK). We found that the cooperative RNAs can sustainably replicate and evolve in microscale water-in-oil droplets.

In the present study, we examined whether a linked RNA could evolve through the long-term replication of the two cooperative RNAs in water-in-oil droplets. We conducted two successive long-term replication experiments by gradually increasing RNA concentrations, since higher RNA concentrations raise the likelihood of generating linked RNAs through recombination or ligation, which could be facilitated by the RNA polymerase [13–15] or occur spontaneously [16–18]. We found that linked RNAs comprising the two cooperating RNAs appeared during the experiment. Sequence analyses revealed that they harbored complete genes, as well as accumulated mutations that were distinct from those in the cooperative RNA fragments. Subsequent biochemical analysis confirmed that an emergent linked RNA replicated the entire sequence by expressing both encoded proteins. Our study provides experimental evidence that supports an evolutionary transition scenario of individually replicating genes to a primitive multi-gene chromosome.

## Results

### Cooperative RNA replication system

The cooperative RNA replication system consists of a reconstituted cell-free translation system derived from *Escherichia coli* [19] and two types of cooperative RNA replicators (Rep- and NDK-RNAs) (Fig 1A) [12]. Rep-RNA (2041 bp) encodes the catalytic subunit of Qβ replicase (*rep* gene), which becomes active upon association with EF-Tu and EF-Ts in the translation system. NDK-RNA (752 bp) encodes NDK (*ndk* gene), which provides a substrate for RNA replication by converting cytidine diphosphate (CDP) into cytidine triphosphate (CTP). Therefore, Rep- and NDK-RNAs cooperatively replicate through the translation of their encoded enzymes. During replication, mutations occur, and occasional recombination generates a parasitic RNA that loses the gene region of either Rep- or NDK-RNA but replicates by exploiting replicase and NDK translated from other RNAs (Fig 1B). We expected that recombination or ligation between Rep- and NDK-RNAs facilitated by Qβ replicase [13–15] would result in the appearance of a linked RNA that harbors both *rep* and *ndk* genes (Fig 1B).

**Fig 1.**
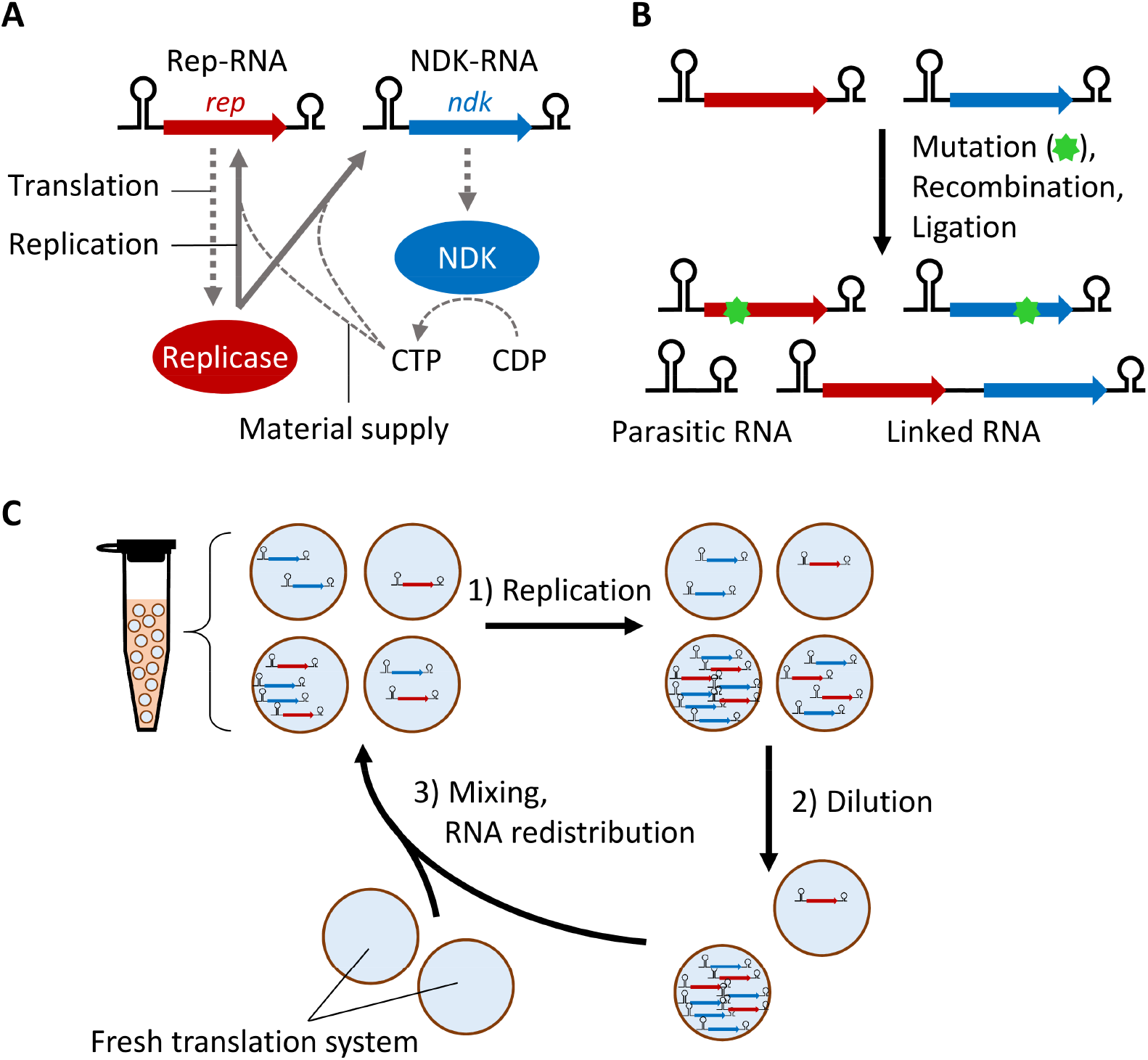
Cooperative RNA replication system. (**A**) Schematic representation of cooperative RNA replication. NDK is translated from NDK-RNA and converts CDP into CTP, a material for RNA replication, whereas replicase is translated from Rep-RNA and replicates both RNAs using the synthesized CTP. (**B**) During replication, mutant Rep- and NDK-RNAs and parasitic RNA that loses gene regions are generated by random mutation and recombination. Similar recombination or ligation could also generate linked RNA that harbors both *rep* and *ndk* genes. (**C**) Schematic representation of long-term replication experiments. 1) Cooperative RNA replication was performed at 37 °C in water-in-oil droplets. 2) Droplets were diluted with new droplets containing a translation system. The dilution rate was typically 5-fold, but 100-fold dilution was used in some rounds. 3) Droplets were vigorously mixed to facilitate their random fusion and division, supplying the translation system to RNA, and RNA was randomly redistributed in the droplets.

### Long-term replication experiments

To investigate whether linked RNAs with the two genes appear from unlinked Rep- and NDK-RNAs, we conducted long-term replication experiments. We repetitively performed (1) RNA replication by incubating the cooperative RNA replication system at 37 °C for 4 or 6 h in water-in-oil droplets, (2) diluted the droplet population typically 5-fold, and (3) induced fusion and division of droplets through vigorous mixing (Fig 1C). In step (3), a fresh translation system was supplied to the RNA population, and the RNA molecules were randomly redistributed among the droplets. We measured Rep- and NDK-RNA concentrations after every replication step using quantitative PCR after reverse transcription (RT-qPCR) with sequence-specific primers. We also measured parasitic RNA concentrations in some rounds using native polyacrylamide gel electrophoresis (PAGE), as their sequences were unknown.

In a previous long-term replication experiment, we maintained RNA concentrations below a certain level (1 nM) by changing the dilution rate every round [12]. This was important because high RNA concentrations cause frequent RNA recombination, producing short parasitic RNA that replicates faster than Rep- and NDK-RNAs. The parasitic RNA propagates throughout the compartments by fusion-division of the compartments and disrupts the cooperation between Rep- and NDK-RNAs [12]. However, in the present study, we increased RNA concentrations to induce recombination and ligation, as these processes produce linked RNA. To achieve a higher RNA concentration while repressing the amplification of parasitic RNA as much as possible, we employed a new protocol to determine the dilution rate; the dilution rate was kept low (5-fold) until RNA concentration reached a particular level, around which parasitic RNAs often appear, and then increased to 100-fold for only a few rounds. We expected that this rapid dilution would circumvent the propagation of parasitic RNAs among the compartments and allow both Rep- and NDK-RNAs to replicate continuously while periodically reaching high concentrations.

In the first long-term replication experiment, we initiated a serial dilution cycle with Rep- and NDK-RNAs obtained in our previous study (hereafter termed e1R0) [12]. In this experiment, we raised the dilution rate from 5-to 100-fold when the RNA concentration reached around 30 nM and successfully continued the replication of both Rep- and NDK-RNAs for 46 rounds (Fig 2A). Parasitic RNA concentrations were measured in four rounds with relatively high Rep- and NDK-RNA concentrations but barely detected (S1A Fig, the detection limit was approximately 10 nM). Next, we evaluated the presence of linked RNA in five rounds with relatively high Rep- and NDK-RNA concentrations by RT-PCR, using primers that could detect RNA with *rep* and *ndk* genes linked in the order 5′-*rep*-*ndk*-3′ (Fig 2B). If the entire region of *rep* and *ndk* genes is retained in the linked RNA, the expected size of the PCR products is between 2136 and 2456 bp. Using agarose gel electrophoresis, we detected approximately 2.5 kbp products in four rounds (Fig 2B), with band intensities roughly correlated with Rep- and NDK-RNA concentrations measured by RT-qPCR (Fig 2C). We also performed RT-PCR using primers for RNA with the two genes linked in the reverse order (5′-*ndk*-*rep*-3′) (S2A Fig) and detected approximately 2.3–2.5 kbp products in only two rounds (S2B Fig), without correlation with Rep- and NDK-RNA concentrations (S2C Fig). These results suggested that linked RNAs appeared during the serial dilution cycle.

**Fig 2.**
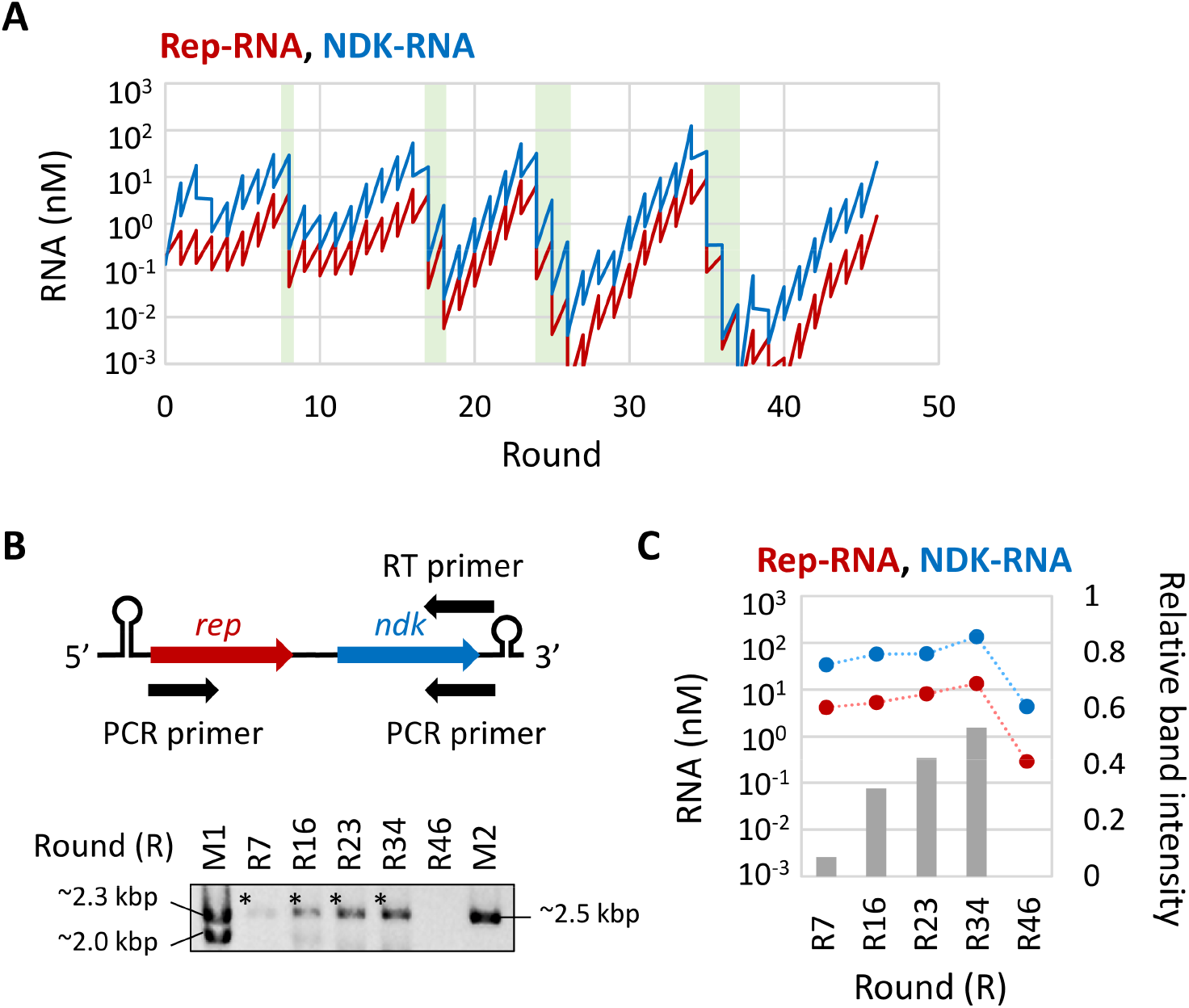
The first long-term replication experiment. (**A**) Changes in Rep-RNA (red) and NDK-RNA (blue) concentrations during the long-term replication experiment, measured by RT-qPCR. The experiment was initiated with Rep- and NDK-RNAs “e1R0” and conducted with high (100-fold) temporal dilution. The replication step was performed at 37 °C for 6 h. Parasitic RNAs were visualized by native PAGE at rounds 8, 17, 24, and 34, but detected only at round 34 (S1A Fig). The parasitic RNA concentration at the round was 33 nM (not plotted). The light green areas highlight the rounds with high dilution. (**B**) RNA samples at the indicated rounds of the long-term replication experiment were subjected to RT-PCR, using primers that could detect 5′-*rep*-*ndk*-3′ (top), and PCR products were analyzed by agarose gel electrophoresis (bottom). M1 and M2 are size markers. Asterisks indicate analyzed bands. (**C**) Relative band intensities of the RT-PCR products to M2 (grey bars, right axis), in comparison with Rep- and NDK-RNA concentrations (red and blue plots, left axis). Dotted lines are plotted for visibility.

Encouraged by the first experiment, we performed a second long-term replication experiment with one of the mutant Rep- and NDK-RNAs obtained at round 46 of the first experiment. These mutant Rep- and NDK-RNAs accumulated six and one mutations, respectively (S1 Table). The replication abilities of the mutant RNA pair (e2R0) were different from those of the original pair (e1R0); Rep- and NDK-RNA replications increased and decreased, respectively (S3 Fig).

In the second long-term replication experiment with the e2R0 pair, we increased the dilution rate from 5-to 100-fold when the RNA concentration reached around 100 nM and successfully continued the replication of both Rep- and NDK-RNAs for 79 rounds (Fig 3A). Parasitic RNA concentrations were measured in rounds 5–7, 20–22, 34–36, 54–56, and 60–62, in which Rep- and NDK-RNA concentrations were relatively high, and detected only in rounds 20–22 (Fig 3A, S1B Fig). The dilution rate was not increased around rounds 35 and 62 when parasitic RNA was undetectable; however, Rep- and NDK-RNA concentrations decreased and then recovered. As a control, we also conducted a continuous replication experiment without changing the dilution rate when the RNA concentration reached 100 nM. In this case, parasitic RNA appeared and replicated excessively (S1C Fig, S4 Fig), and then the concentrations of Rep- and NDK-RNAs decreased to undetectable levels, confirming the importance of changing the dilution rate.

**Fig 3.**
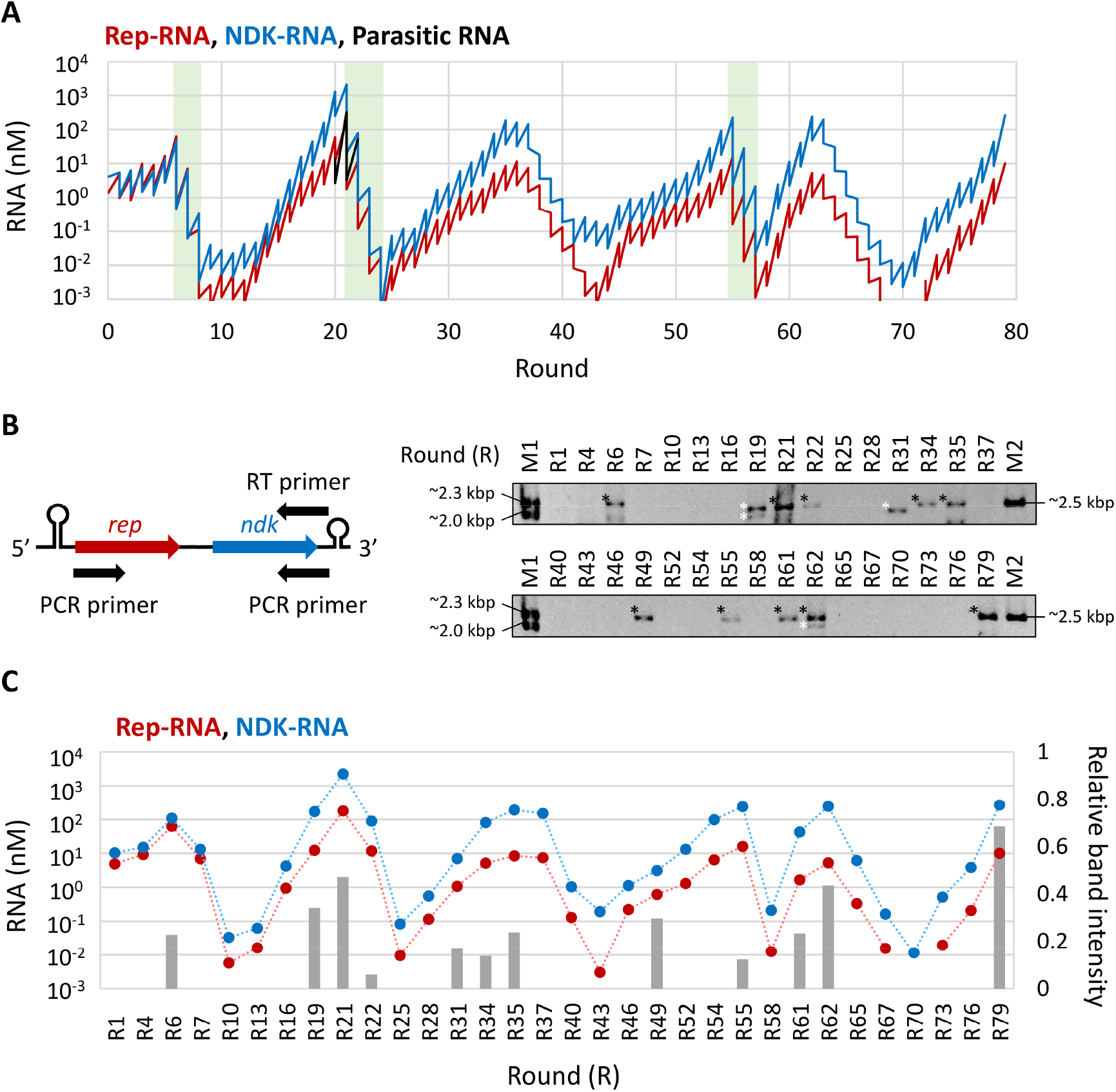
The second long-term replication experiment. (**A**) Changes in Rep-RNA (red), NDK-RNA (blue), and parasitic RNA (black) concentrations during the long-term replication experiment. The experiment was initiated with Rep- and NDK-RNAs “e2R0” and conducted with high (100-fold) temporal dilution. The replication step was performed at 37 °C for 4 h. RNA concentrations were measured by RT-qPCR (Rep- and NDK-RNAs) or based on native PAGE (parasitic RNAs, S1B Fig). Detection of parasitic RNAs was attempted at the rounds described in the main text. Rounds with high dilution are highlighted in light green. (**B**) Detection of putative linked RNAs that harbor both *rep* and *ndk* genes (5′-*rep*-*ndk*-3′) in the RNA samples in every 1–3 rounds of the long-term replication experiment. RT-PCR was performed using the indicated primers (left), and PCR products were analyzed by agarose gel electrophoresis (right). M1 and M2 are size markers. Asterisks indicate analyzed bands; white ones indicate shorter sizes. (**C**) Relative band intensities of RT-PCR products to M2 (grey bars, right axis), in comparison with Rep- and NDK-RNA concentrations (red and blue plots, left axis). Dotted lines are plotted for visibility.

Next, we examined the appearance of a linked RNA by RT-PCR every 1–3 rounds, using primers that could detect 5′-*rep*-*ndk*-3′ (Fig 3B). We detected approximately 2.3–2.5 kbp PCR products at rounds 6, 21, 22, 34, 35, 49, 55, 61, 62, and 79 and seemingly smaller (2.0–2.3 kbps) products at rounds 19, 31, and 62. The relative intensities of the bands, if detected, were as strong as those in the first long-term replication experiment (Fig 2B) throughout the rounds. The band intensities were strong in rounds with high Rep- and NDK-RNA concentrations (Fig 3C), with some exceptions (e.g., rounds 22, 37, 54, and 55) where the product bands were undetected or barely detected despite high Rep- and NDK-RNA concentrations. The concentration of a potentially linked RNA in the population at round 79 was estimated to be less than 1 nM, which is less than 10% of either of the unlinked RNAs, by RT-qPCR using primers that could detect the linkage region of 5′-*rep*-*ndk*-3′. We also performed RT-PCR at rounds 6, 19, 34, 54, 62, and 79 using primers for 5′-*ndk*-*rep*-3′ (S2D Fig, S2E Fig); we detected approximately 2.3–2.5 kbps products at only round 62.

### Sequence analysis

Next, we obtained 16 clones of the potentially linked RNA products (5′-*rep*-*ndk*-3′) at round 79 of the second long-term replication experiment and analyzed their sequences. All analyzed clones harbored the entire region of both *rep* and *ndk* genes in the 5′-*rep*-*ndk*-3′ order, confirming that the obtained long RNAs were the linked products of Rep- and NDK-RNAs (hereafter referred to as RepNDK-RNA). We found two major ways by which Rep- and NDK-RNAs were linked. In 56% of the RepNDK-RNA clones, Rep- and NDK-RNAs were simply connected end-to-end without deletion or insertion. In 38% of the clones, a single C was inserted between the connected Rep- and NDK-RNAs (Fig 4A, linkage region). At the NDK-RNA site around the linkage, we also detected G3A and U12G in 75% and 88% of the RepNDK-RNA clones, respectively.

**Fig 4.**
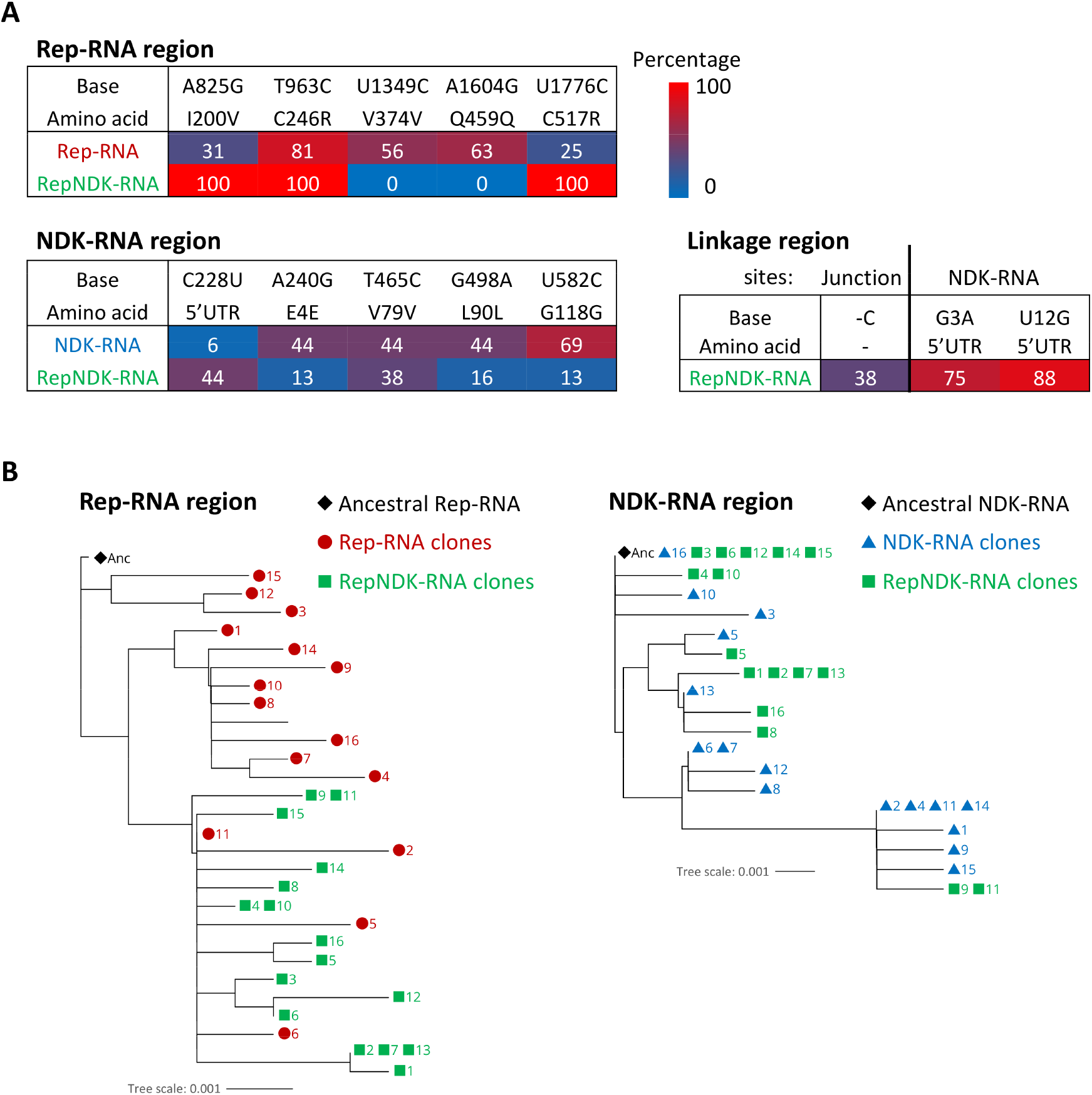
Accumulated mutations in the linked and unlinked RNAs. (**A**) Mutations detected in more than 30% of the 16 clones of any RNA species analyzed at round 79 are listed with their percentages. The sequence of the RepNDK-RNA clones was divided into Rep-RNA, NDK-RNA, and linkage regions for comparison with the Rep- and NDK-RNA clones. All detected mutations are listed in S2 Table and S3 Table. (**B**) Phylogenetic trees for each Rep-(left) and NDK-(right) RNA region were constructed from all the analyzed clones. Leaves representing different RNA species (Rep-, NDK-, or RepNDK-RNA) are marked with different colored symbols, followed by clone numbers. The ancestral Rep- and NDK-RNAs (Anc) were analyzed and displayed together.

Next, to investigate the origin of RepNDK-RNA at round 79, we compared the sequences of the RepNDK-RNA clones with those of unlinked Rep- and NDK-RNAs. If RepNDK-RNA was continuously replicated during the long-term replication experiment, they could have accumulated mutations different from those found in unlinked Rep- and NDK-RNAs in the same round. However, if RepNDK-RNA appeared from Rep- and NDK-RNAs around round 79 or just as artificial products during the RT-PCR process, they should have a similar set of mutations. To distinguish these possibilities, we obtained 16 clones of unlinked Rep- and NDK-RNAs at round 79 and analyzed their sequences. The Rep- and NDK-RNA clones contained 3–8 and 0–5 mutations, respectively (S2 Table, S3 Table).

Fig 4A lists all mutations detected in more than 30% of the clones of any of the three RNA species (unlinked Rep- and NDK-RNAs and linked RepNDK-RNAs). The three types of RNAs accumulated different mutations at their corresponding sites. For example, A825G and U1776C were found in 31% and 25% of the Rep-RNA clones, respectively, whereas both mutations were detected in 100% of the RepNDK-RNA clones. In contrast, U1349C and A1604G were observed in 56% and 63% of the Rep-RNA clones, respectively, but were not detected in the RepNDK-RNA clones. Similarly, C228U was found in only 6% of the NDK-RNA clones but in 44% of the RepNDK-RNA clones, whereas U582C was detected in 69% of the NDK-RNA clones but in only 13% of the RepNDK-RNA clones. These different mutation patterns indicate that the RepNDK-RNAs analyzed at round 79 were not derived from Rep- and NDK-RNAs of the same round. These results also suggested that RepNDK-RNA was maintained in the population for a certain period, long enough to form its own lineage.

We further illustrated the evolutionary relationships between Rep- or NDK-RNA and RepNDK-RNA using phylogenetic trees for each of the Rep- and NDK-RNA regions (Fig 4B). The three RNA species are represented by different symbols. For the Rep-RNA region, most of the unlinked Rep-RNA clones (red circles) and linked RepNDK-RNA clones (green squares) were clustered in different branches, while some Rep-RNA clones (clones 11, 2, 5, and 6) were present in the RepNDK-RNA cluster. For the NDK-RNA region, the unlinked NDK-RNA clones (blue triangles) and linked RepNDK-RNA clones (green squares) were mixed in the tree. These results suggested that RepNDK-RNA formed lineages that were not completely but partially distinct from the unlinked RNAs.

### Biochemical characterization of RepNDK-RNA

To understand the biochemical characteristics of the linked RepNDK-RNA at round 79, we selected a clone containing only the most common mutation set (A825G, T963C, U1776C, and -2042C in the Rep-RNA region and G3A and U12G in the NDK-RNA region), which was found in 25% of the 16 clones. For comparison, we also chose a pair of unlinked Rep- and NDK-RNAs containing only the most common mutation sets at round 79. The selected Rep-RNA contained U963C, U1349C, and A1604G, found in 44% of the Rep-RNA clones, whereas the selected NDK-RNA contained A240G, U465C, G498A, and U582C, found in 38% of the NDK-RNA clones. The replication of these Rep- and NDK-RNAs (e2R79) was similar to that of the ancestral Rep- and NDK-RNA pair (e2R0) with slightly improved NDK-RNA replication (S3 Fig).

RepNDK-RNA was expected to replicate by expressing both encoded replicase and NDK. To test this, we incubated the RepNDK-RNA clone in a reconstituted translation system containing fluorescent-labeled lysyl-tRNA. The newly synthesized proteins were analyzed by sodium dodecyl sulfate (SDS)-PAGE, followed by fluorescent imaging. We found that bands corresponding to both the replicase subunit and NDK were detected for the RepNDK-RNA (Fig 5A). The protein amounts estimated from the band intensities were approximately half of those of the Rep- and NDK-RNAs (Fig 5B). Next, we investigated whether the translation of the two proteins was coupled with RepNDK-RNA replication. During RNA replication, the replicase first synthesizes the minus (complementary) strand, which is then recognized by the replicase for plus-strand synthesis (Fig 5C). We incubated the RepNDK-RNA clone with a translation system in water-in-oil droplets at 37 °C for 4 h and analyzed the synthesis of full-length plus and minus strands by RT-PCR. Agarose gel electrophoresis showed that the intensities of bands corresponding to both strands increased after incubation (Fig 5D), demonstrating the ability of the RepNDK-RNA to replicate without the help of unlinked RNAs. To quantitatively compare the replication of the linked RNA with that of the unlinked RNAs, we measured the replication of the RepNDK-RNA by RT-qPCR. The extent of RepNDK-RNA replication was less than 10% of Rep- and NDK-RNA replications (Fig 5E), which may be caused by the reduced translation activity.

**Fig 5.**
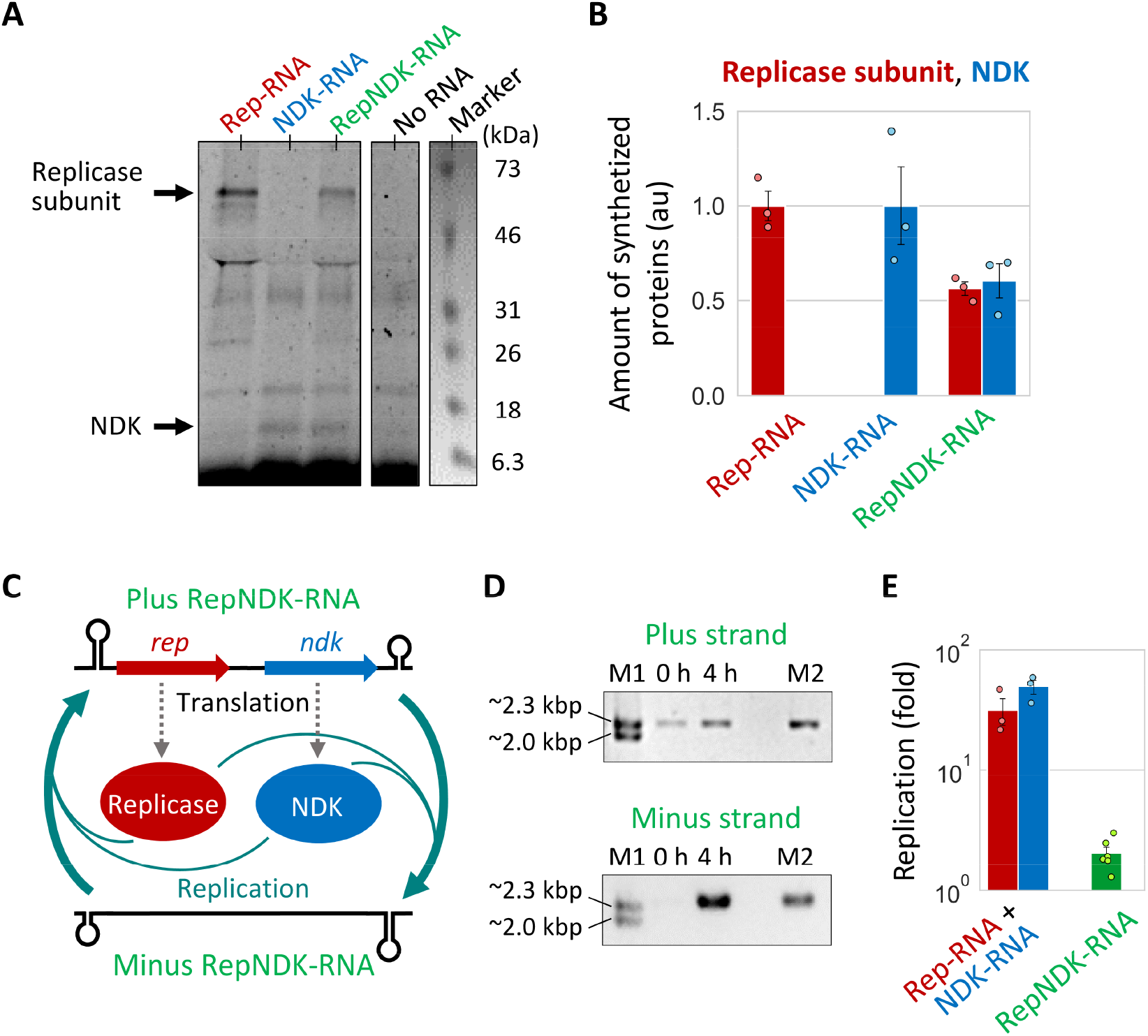
Biochemical properties of the RNA clones at round 79. (**A**) SDS-PAGE of the translation products after incubation of the Rep-, NDK-, and RepNDK-RNA clones (300 nM) at 37 °C for 12 h in a modified translation system containing a fluorescently labeled lysine tRNA. Neither CTP nor CDP was present to prevent RNA replication. An analyzed fluorescent gel image is shown, in parallel to a trimmed white-light image of the same gel to visualize a pre-stained molecular weight marker. The expected bands of the replicase subunit (~64 kDa) and NDK (~15 kDa) are indicated by the black arrows. (**B**) The amounts of synthesized proteins, normalized to those of the Rep- or NDK-RNA clones (replicase subunit or NDK, respectively). Error bars indicate standard errors (n=3). (**C**) The expected replication scheme of RepNDK-RNA. The translation of the two encoded proteins induced RNA replication, completed with minus strand (complementary strand) synthesis from a plus strand (template) and following plus strand synthesis from the minus strand. (**D**) The RepNDK-RNA clone (10 nM) was incubated with the translation system in water-in-oil droplets at 37 °C for 4 h, and the replications of entire plus and minus strands were analyzed by RT-PCR and agarose gel electrophoresis. M1, size marker. M2, control PCR product of the RepNDK-RNA clone. (**E**) The replication amount of the RepNDK-RNA clone was measured by RT-qPCR in comparison with that of the Rep- and NDK-RNA pair (10 nM each). Error bars indicate standard errors (n=3–6).

## Discussion

The integration of distinct genetic information on a primitive chromosome has been considered a major evolutionary transition in the development of life [2–4]. In this study, we demonstrated that such a linkage emerged during the long-term replication of two cooperative RNAs encoding replication and metabolic enzymes in cell-like compartments.

The linked RNAs that existed in the final population retained both genes, maintained their translation and replication abilities, and accumulated mutations different from those found in the unlinked RNAs (Fig 4 and 5), suggesting that the linked RNA continuously replicated in the RNA population for at least a certain period. These results provide experimental evidence that individual RNA replicators encoding different genes can assemble into a single RNA molecule through Darwinian evolution. This process may explain the origin of primitive chromosomes during prebiotic evolution.

Although the accumulation of unique mutations in RepNDK-RNA indicated their maintenance in the population, at round 79, the linked RNA replicated less efficiently than the pair of Rep- and NDK-RNAs (Fig 5E), possibly because of its lower translational activity (Fig 5B), extended length, and reorganized RNA structures. How the linked RNA was maintained in the population while competing with the unlinked RNAs that replicate faster remains unexplained. One possible reason could be that, unlike unlinked Rep- and NDK-RNAs, RepNDK-RNA can always replicate on its own after redistribution in the droplet population, as suggested by theoretical studies [8,9,11]. This effect is expectedly enhanced when the stochastic mis-encapsulation of Rep- and NDK-RNAs becomes effective at low RNA concentrations [12]. Experiments using lower RNA concentrations for long-term replication could provide more selective advantages for RepNDK-RNA.

The generation of a linkage between RNA fragments is not unique to our cooperative RNA replication system. Diverse RNA-dependent RNA polymerases (RdRp), including Qβ replicase, are known to cause intermolecular RNA recombination during replication [20–22], which can link multiple RNA molecules. End-to-end ligation, as detected in the present study (with or without single nucleotide insertion), has also been observed for various RdRp such as through end-to-end template switching [14,23]. Thus, whether linked RNAs can evolve in other *in vitro* RNA replication systems using different RNA replicases [24] needs to be investigated. Furthermore, even before the emergence of proteins, such recombination and ligation of RNA molecules could have occurred spontaneously [16–18] or with the assistance of catalytic RNAs [25–29]. Therefore, it is reasonable to assume that the linkage of cooperative RNA molecules was prevalent during the early evolution of life. We also developed a new method that allows the long-term replication of cooperative RNAs, even if the RNA concentrations increase tentatively and parasitic RNA emerges. Our previous study suggested that maintaining a relatively low RNA concentration was crucial for sustained replication of the cooperative system, and continuous RNA replication was achieved only by controlling the dilution rate at every replication round [12]. However, the present study demonstrated that sustainable cooperative RNA replication does not require strict regulation of RNA concentration, as previously believed, supporting the robustness of molecular cooperation. A higher RNA concentration also increases the chance of recombination or ligation of RNAs and may facilitate the evolution of a primitive chromosome.

The evolution of a linked RNA from multiple RNAs that encode different genes may also be useful for developing an artificial replicable RNA genome. To the best of our knowledge, RepNDK-RNA is the first artificial RNA genome that replicates based on the translation of more than one encoded protein. The manual expansion of an RNA genome by introducing new genes is challenging, as it typically disrupts RNA structures and impairs its ability to undergo replication and translation. Our previous study connected two RNAs based on the predicted RNA structures of linked RNAs while maintaining their replication coupled with the translation of one encoded protein [30]. However, further introduction of genes would make genome expansion more difficult as the accuracy of RNA structural production decreases with increasing RNA length. Although further refinement is necessary, the evolutionary technique to integrate genes into a single RNA genome, as demonstrated here, could be utilized to create an optimized RNA genome encoding multiple proteins and contribute to the development of artificial cells that possess a genome replication system [31,32].

## Materials and Methods

### Plasmids and RNAs

Two plasmids, each encoding Rep- or NDK-RNA “e1R0,” were obtained in the previous study as the plasmids encoding Rep- or NDK-RNA “Evo” [12]. The other plasmids, each encoding one of the two Rep- and NDK-RNA clones (“e2R0” or “e2R79”) or the RepNDK- RNA clone, were obtained in the present study through cloning, as described below, and site-specific mutagenesis. For the RepNDK-RNA clone, the entire cDNA length was PCR-amplified from the corresponding plasmid. The cDNA of the RepNDK-RNA clone and the plasmids for all other RNA clones were subjected to digestion with Sma I (Takara) and *in vitro* transcription with T7 RNA polymerase (Takara). All transcribed RNAs were purified using the RNeasy Mini Kit (QIAGEN).

### Preparation of the reconstituted translation system

The composition of the reconstituted translation system was as described previously [12] (based on the reconstituted *Escherichia coli* translation system [19]) except that the trigger factor was omitted because it was not essential for translation and contained a high level of NDK activity that could not be reduced by the following purification steps. All protein components of the translation system were purified by two successive affinity column chromatography in a stringent buffer to further reduce the remaining NDK activity derived from *Escherichia coli*. For all proteins except ribosomes, the re-purification procedure was the same as that used for removing tRNA from EF-Tu, described in our previous study [33]. Ribosomes were purified as described previously [19] and then washed with another stringent buffer (20 mM Hepes-KOH (pH 7.6), 6 mM magnesium acetate, 7 mM 2-mercaptoethanol, 1% Triton X-100, 1 mM dithiothreitol, and 0.33 M potassium chloride).

Briefly, the ribosomes were diluted 20-fold with the stringent buffer and ultracentrifuged at 150,000 g for 2 h at 4 °C. After removing the supernatant, the precipitate was rinsed with 70S buffer (20 mM Hepes-KOH (pH 7.6), 6 mM magnesium acetate, 7 mM 2-mercaptoethanol, and 0.03 M potassium chloride). After carefully removing the remaining buffer, the precipitate was dissolved in 70S buffer. Then, the solution was diluted again with the stringent buffer and collected by ultracentrifugation, as described above. To remove the residual stringent buffer, the final ribosome solution was diluted with 70S buffer 10-fold and then concentrated using Amicon Ultra (30 kDa cut, Merck) three times.

### Assay of remaining NDK activity

The remaining NDK activity in each protein component before and after the purification step was estimated as the activity to convert adenosine diphosphate (ADP) into adenosine triphosphate (ATP). First, each protein component (twice the concentration of that in the translation system) was incubated with 2.5 mM ADP and 1.25 mM CTP in a reaction buffer (100 mM Hepes-KOH (pH 7.6), 70 mM glutamic acid potassium salt, 0.375 mM spermidine, and 11 mM magnesium acetate) at 37 °C for 2 h. Then, an aliquot of the reaction was used to determine the amount of synthesized ATP using ATP Assay Kit (Colorimetric/Fluorometric) (Abcam). The assay was performed at 37 °C, and the fluorescence intensity was measured using Mx3005P Real-Time PCR System (Agilent Technologies) as the indicator of ATP synthesis (S5 Fig).

### Long-term replication experiment

The experiments were performed as described previously [12] with several modifications. The translation system (10 μL) containing certain concentrations of Rep- and NDK-RNAs “e1R0” (Fig 2A) or “e2R0” (Fig 3A and S4 Fig) were added to 1,000 μL of buffer-saturated oil and mixed vigorously with a homogenizer (POLYTRON PT 1300D, KINEMATICA) at 16,000 rpm for 1 min on ice to obtain water-in-oil droplets. The preparation of the saturated oil was described in the previous study [34]. The droplets were incubated at 37 °C for 6 h (Fig 2A) or 4 h (Fig 3A and S4 Fig) to induce RNA replication through protein translation. After incubation, an aliquot of the droplets was diluted 5-fold or 100-fold with fresh buffer-saturated oil, as described in the main text and shown in the corresponding figures. The solution was mixed with 10 μL of the translation system and homogenized using the same method to obtain a new droplet population, followed by incubation at 37 °C for 4 or 6 h for the next round of RNA replication. In Fig 2A, the original droplet population was diluted 100-fold using the same method as that before incubation to make the initial population (approximately 0.1 nM of Rep- and NDK-RNAs). Rep- and NDK-RNA concentrations were determined at every round by RT-qPCR with primers 1 and 2 (Rep-RNA) or 3 and 4 (NDK-RNA) (S4 Table). The measurement was performed after diluting the droplets 100-fold with 1 mM EDTA (pH 8.0) and using One Step TB Green PrimeScript PLUS RT-PCR Kit (Takara).

### Measurement of parasitic RNA concentration

In some rounds of the long-term replication experiments, the water phase containing RNA was collected from the droplets (100 μL) by centrifugation (22,000 g, 5 min). The recovered phase was mixed with diethyl ether (40 μL) and centrifuged (11,000 g, 1 min) to remove the diethyl ether phase. Then, RNA was purified with the RNeasy Mini Kit (QIAGEN) and subjected to 8% polyacrylamide gel electrophoresis in 1×TBE buffer. The fluorescence intensities of bands corresponding to parasitic RNAs were quantified using ImageJ (NIH) after staining with SYBR Green II (Takara). The concentrations of parasitic RNAs were determined from the intensities based on a dilution series of a standard parasitic RNA (s222 [35]).

### RT-PCR and Sequence analysis

RNA samples were obtained from the long-term replication experiments as described above. Rep- and NDK-RNAs were reverse transcribed with primer 8, PCR-amplified with primers 7 and 8 (S4 Table), and separated using 0.8% agarose gel electrophoresis with E-Gel CloneWell (Thermo Fisher Scientific). The cDNA samples were cloned into a pUC19 vector (PCR-amplified with primers 11 and 12) using In-Fusion HD Cloning Kit (Takara). After transformation into *Escherichia coli*, the sequences of randomly selected plasmids were analyzed. Similarly, RepNDK-RNA was reverse transcribed with primer 10, PCR-amplified with primers 9 and 10, size-selected, and cloned using the same method. The 5′ and 3′ untranslated regions of the RepNDK-RNA clones could not be retrieved to distinguish them from Rep- and NDK-RNAs in the same RNA population. The sequence information of the linkage region was obtained only for the RepNDK-RNA clones. To detect putative elongated RNAs with *rep* and *ndk* genes linked in this order (Fig 2C and 3C) or the reverse order (S2 Fig), RT-PCR was performed with primers 9 and 10 or 13 and 14, respectively. PCR products were visualized by agarose gel electrophoresis and stained with SAFELOOK Green Nucleic Acid Stain (FUJIFILM), and band intensities were determined using ImageJ (NIH).

### Phylogenetic analysis

The sequences of the Rep-, NDK-, and RepNDK-RNA clones obtained at round 79 of the second long-term replication experiment (Fig 3A) were subjected to phylogenetic analysis. The RepNDK-RNA clones were divided into Rep-RNA and NDK-RNA regions and compared with the Rep- and NDK-RNA clones. Only the sequence regions of the RepNDK-RNA clones and their respective unlinked RNAs for which mutation information was commonly available were used for the analysis. Phylogenetic trees were created using the neighbor-joining method in MEGA11 [36] by assuming the same mutation rate for point mutations, single nucleotide deletions, and single nucleotide insertions. Phylogenetic trees were visualized using Interactive Tree Of Life (iTOL) [37].

### Translation-coupled RNA replication experiments

A pair of Rep- and NDK-RNA clones (10 nM each) or the RepNDK-RNA clone (10 nM) was incubated with the translation system in water-in-oil droplets at 37 °C for 4 h. The concentration of each RNA clone was determined by RT-qPCR as described above. Primers 5 and 6 were used for the RepNDK-RNA clone (S4 Table).

### Analysis of protein translation

300 nM of Rep-, NDK-, or RepNDK-RNA was incubated at 37 °C for 12 h in a translation system and FluoroTect GreenLys tRNA (Promega), with neither CTP nor CDP to preclude RNA replication. In this experiment, we used a translation system before excessive purification of protein components, as described previously [38], to enhance translation efficiency and obtain measurable amounts of proteins. After translation, an aliquot was treated with 0.1 mg/mL RNase A (QIAGEN) at 37 °C for 15 min to digest fluorescently labeled lysine tRNA. Then, the solution was incubated at 95 °C for 4 min in SDS sample buffer (50 mM tris(hydroxymethyl)aminomethane hydrochloride (Tris-HCl, pH 7.4), 2% SDS, 0.86 M 2-mercaptoethanol, and 10% glycerol) and subjected to SDS-PAGE using a 10–20% gradient gel (Funakoshi, Japan). The synthesized proteins that incorporated fluorescently labeled lysine were visualized using FUSION-SL4 (Vilber-Lourmat), and band intensities were determined using ImageJ (NIH).

## Acknowledgments

We are grateful to Ms. Nana Kuroda for technical support.

## Supporting information captions

**S1 Fig. Detection of parasitic RNAs during the long-term replication experiments**.

**S2 Fig. Detection of putative linked RNAs harboring *rep* and *ndk* genes in the reverse order**.

**S3 Fig. Translation-coupled cooperative RNA replication experiment**.

**S4 Fig. Long-term replication experiment without high temporal dilution**.

**S5 Fig. Contamination levels of NDK in the translation system**.

**S1 Table. The list of mutations in the Rep- and NDK-RNA clones obtained at round 46 of the long-term replication experiment shown in Fig 2A**.

**S2 Table. The list of mutations in the Rep- and RepNDK-RNA clones obtained at round 79 of the long-term replication experiment shown in Fig 3A**.

**S3 Table. The list of mutations in the NDK- and RepNDK-RNA clones obtained at round 79 of the long-term replication experiment shown in Fig 3A**.

**S4 Table The list of primers (from 5′ end to 3′end)**.

## Author Contributions

**Conceptualization:** Kensuke Ueda, Ryo Mizuuchi, Norikazu Ichihashi.

**Data curation:** Kensuke Ueda.

**Formal analysis:** Kensuke Ueda, Ryo Mizuuchi.

**Funding acquisition:** Ryo Mizuuchi, Norikazu Ichihashi.

**Investigation:** Kensuke Ueda, Ryo Mizuuchi, Norikazu Ichihashi.

**Methodology:** Kensuke Ueda, Ryo Mizuuchi, Norikazu Ichihashi.

**Project Administration:** Ryo Mizuuchi, Norikazu Ichihashi.

**Supervision:** Ryo Mizuuchi, Norikazu Ichihashi.

**Validation:** Kensuke Ueda, Ryo Mizuuchi, Norikazu Ichihashi.

**Visualization:** Kensuke Ueda, Ryo Mizuuchi.

**Writing – original draft:** Kensuke Ueda, Ryo Mizuuchi.

**Writing – review** & **editing:** Kensuke Ueda, Ryo Mizuuchi, Norikazu Ichihashi.

