## Supplementary Information for "Emergence of linkage between cooperative RNA replicators encoding replication and metabolic enzymes thorough experimental evolution"

**This file includes:**

Figs. S1 to S5

Tables S1 to S4

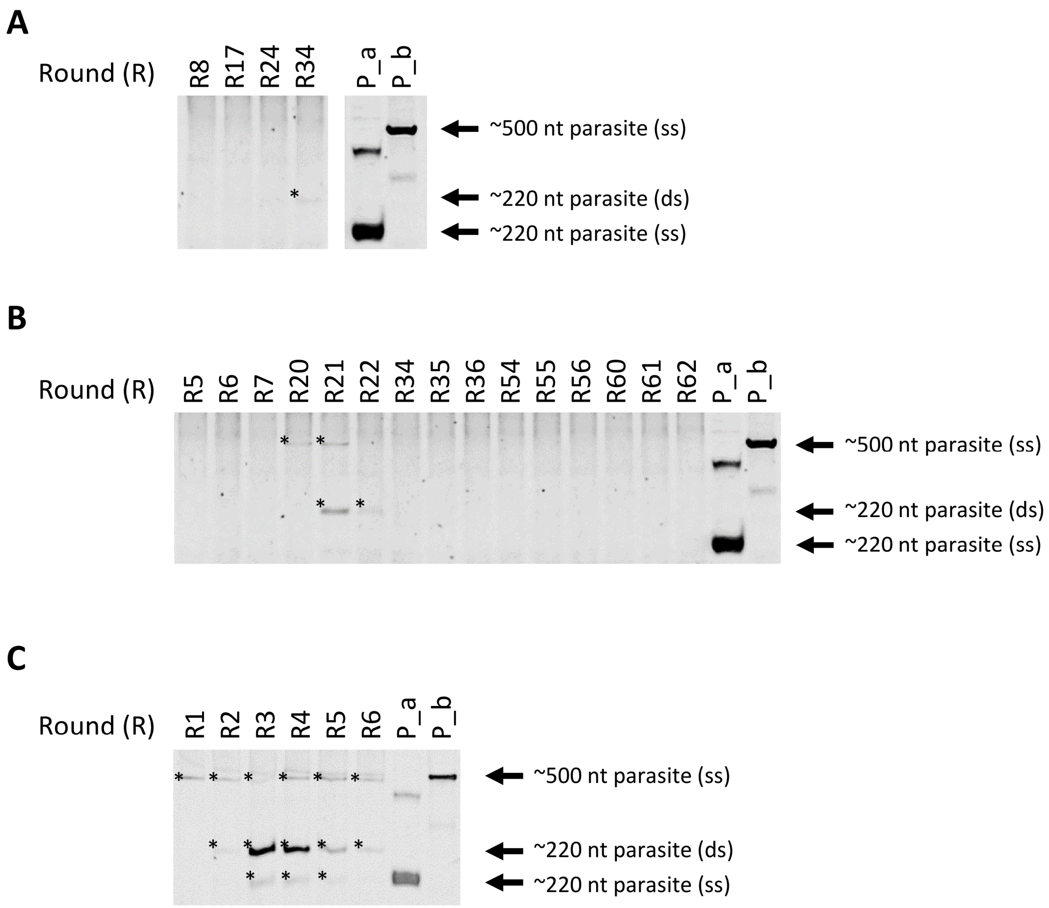


**S1 Fig. Detection of parasitic RNAs during the long-term replication experiments.** (**A, B, C**) Native PAGE of RNA mixtures during the long-term replication experiments shown in Fig 2A (A), Fig 3A (B), and S4 Fig (C). P_a and P_b are controls of commonly appearing parasitic RNAs of known sizes. Asterisks indicate the bands whose intensities were quantified. The expected parasitic RNA bands and their sizes are shown on the right. ss, single-strand. ds, double-strand.

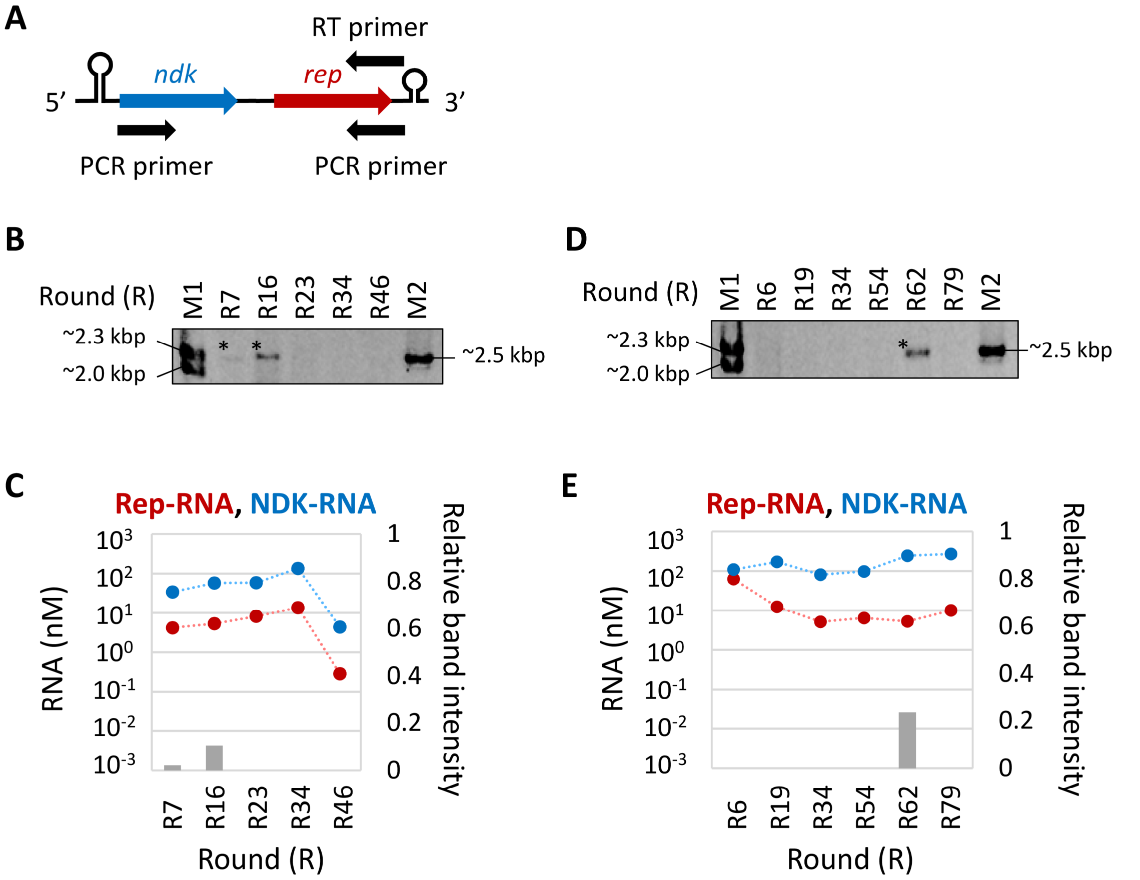


**S2 Fig. Detection of putative linked RNAs harboring *rep* and *ndk* genes in the reverse order.** (**A**) RNA samples in the long-term replication experiments were subjected to RT-PCR using the primers that could detect 5′-*ndk*-*rep*-3′. (**B**) The PCR products were analyzed by agarose gel electrophoresis for RNA samples of the first long-term replication experiment (Fig 2A). M1 and M2 are size markers. Asterisks indicate analyzed bands. (**C**) Relative band intensities of the RT-PCR products to M2 (gray bars, right axis), in comparison with Rep- and NDK-RNA concentrations (red and blue plots, left axis). Dotted lines are plotted for visibility. (**D**, **E**) The same analyses were performed for the RNA samples in the second long-term replication experiment (Fig 3A).

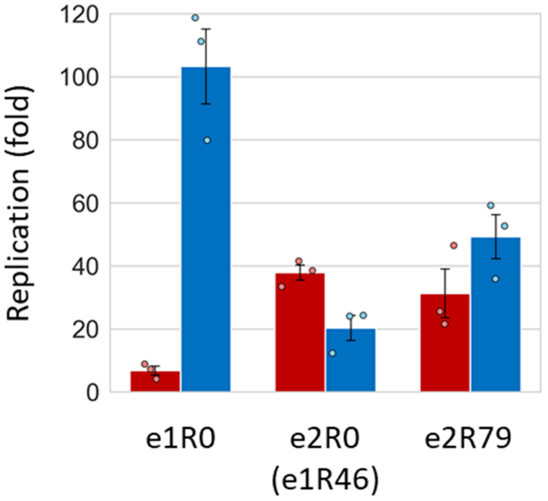


**S3 Fig. Translation-coupled cooperative RNA replication experiment.** A pair of Rep- and NDK-RNAs (10 nM each) was incubated with a translation system in water-in-oil droplets at 37°C for 4 h, and their replication was measured by RT-qPCR. e1R0, e2R0 (e1R46), and e2R79 represent the ancestral RNA clones in the first long-term replication experiment (Fig 2A), the ancestral clones in the second long-term replication experiment (Fig 3A) (obtained at round 46 of the first experiment), and the clones obtained at round 79 of the second experiment, respectively. Error bars indicate standard errors (n=3)

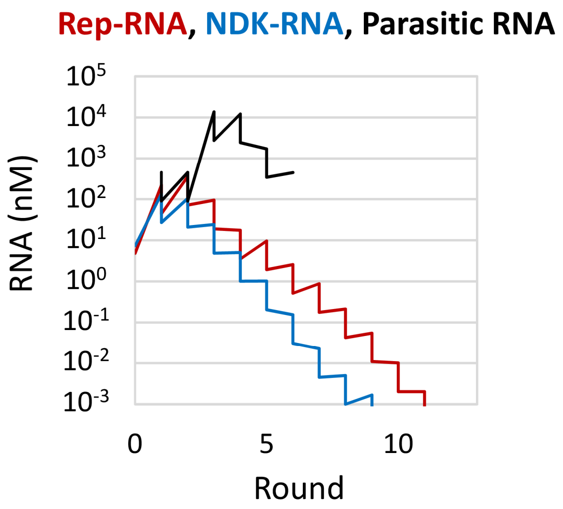


**S4 Fig. Long-term replication experiment without high temporal dilution.** Changes in Rep-RNA (red), NDK-RNA (blue), and parasitic RNA (black) concentrations during the long-term replication experiment. The experiment was initiated with Rep- and NDK-RNAs "e2R0" and conducted without high temporal dilution. The replication step was performed at 37°C for 4 h. RNA concentrations were measured by RT-qPCR (Rep- and NDK-RNAs) or based on native PAGE (parasitic RNAs, S1C Fig).

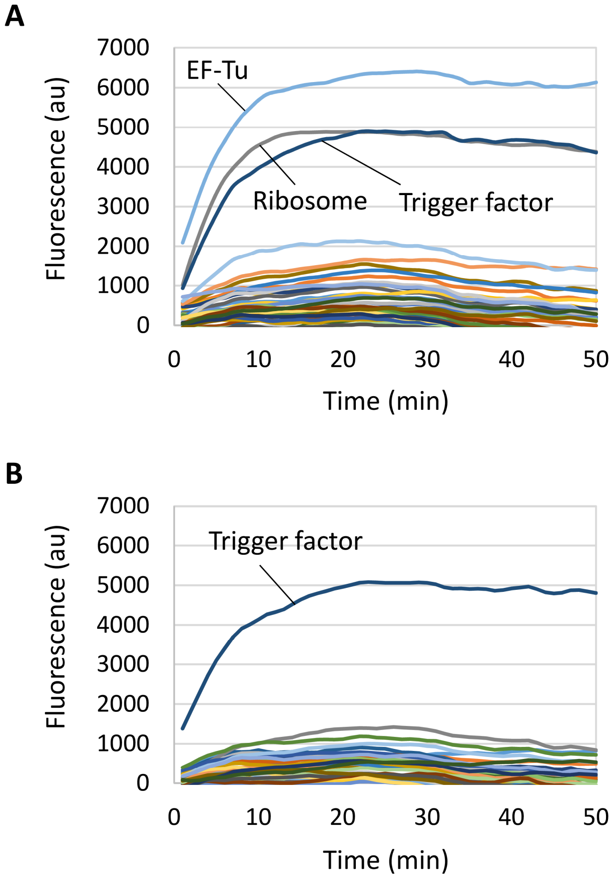


**S5 Fig. Contamination levels of NDK in the translation system.** (**A, B**) The contamination level of NDK in each protein component before (A) and after (B) re-purification was measured as the increase in fluorescence using the ATP assay (see Materials and Methods). The protein names with high levels of contamination are indicated.

**S1 Table. The list of mutations in the Rep- and NDK-RNA clones obtained at round 46 of the long-term replication experiment shown in Fig 2A.**

| | | Rep-RNA clones | | | | | |
|---|---|---|---|---|---|---|---|
| Base | Amino acid | 1 | 2 | 3 | 4 | 5 | 6 |
| A49G | 5'UTR | | | | | | + |
| A290G | T21T | | | + | | | |
| C467U | D80D | | + | | | | |
| U504C | W93R | | | | + | | |
| G697A | G157D | | | | + | | |
| C766U | T180M | + | | | | | |
| U830C | S201S | | | | | + | |
| A982G | N252A | | | + | | | |
| G1059U | A278S | | | | + | | |
| U1157C | P310P | | | | | | + |
| U1199C | G324G | + | | | | | |
| U1235C | F336F | | | + | | | |
| U1278C | S351P | + | | + | | + | + |
| U1392C | S389P | | | | | + | |
| U1739C | A504A | + | | + | | + | + |
| A1792G | N522S | + | | + | | + | + |
| A1848G | I541V | + | | + | | + | + |

| | | NDK-RNA clones | | | | | | | | | |
|---|---|---|---|---|---|---|---|---|---|---|---|
| Base | Amino acid | 1 | 2 | 3 | 4 | 5 | 6 | 7 | 8 | 9 | 10 |
| A53G | 5'UTR | | | + | | | | | + | | |
| C122U | 5'UTR | | | | | | | + | | | |
| U199C | 5'UTR | | | | | | | + | | | |
| U230C | START1T | | | | | + | | | | | |
| G232A | A2T | | + | | | | | | | | |
| A264G | P12P | | + | | | | | | | | |
| U272C | V15A | | | + | | | | | + | | |
| A281G | N18S | | + | | | | | | | | |
| G290U | G21V | | | | | | + | | | | |
| U306- | shift | | | | | + | | | | | |
| A396G | G57G | | | | | + | | | | | |
| U414C | G62G | | | + | | + | | | + | | |
| A505G | T93A | | | | | | + | | | | |
| U643C | C139R | | | | + | | | | | | |
| U658C | STOP144Q | | | + | | | | | + | | |
| U675C | 3'UTR | | + | | | | | | | + | + |

The highlighted clones (Rep-RNA clone 6 and NDK-RNA clone 10) were used for further long-term replication experiments (Fig 3A and S4 Fig).

**S2 Table. The list of mutations in the Rep- and RepNDK-RNA clones obtained at round 79 of the long-term replication experiment shown in Fig 3A.**

| | | Rep-RNA clones | | | | | | | | | | | | | | | | RepNDK-RNA clones | | | | | | | | | | | | | | | |
|---|---|---|---|---|---|---|---|---|---|---|---|---|---|---|---|---|---|---|---|---|---|---|---|---|---|---|---|---|---|---|---|---|---|
| | | 1 | 2 | 3 | 4 | 5 | 6 | 7 | 8 | 9 | 10 | 11 | 12 | 13 | 14 | 15 | 16 | 1 | 2 | 3 | 4 | 5 | 6 | 7 | 8 | 9 | 10 | 11 | 12 | 13 | 14 | 15 | 16 |
| G40A | | + | | | | | | | | | | | | | | | | | | | | | | | | | | | | | | | |
| A46G | | + | | | | | | | | | | | | | | | | | | | | | | | | | | | | | | | |
| G49A | | + | | | | | | | | | | | | | | | | | | | | | | | | | | | | | | | |
| U204C | | | | | | | | | | | | | | | | | | Not analyzed | | | | | | | | | | | | | | | |
| C207- | | | | | | | | | | | | | | | | + | | | | | | | | | | | | | | | | | |
| U208- | | | | | | | | | | | | | | | | + | | | | | | | | | | | | | | | | | |
| A269G | G14G | | | | | | | | | | | | | | + | | | | | | | | | | | | | | | | | | |
| C273U | R16C | | | | | | | | | | | | | | | | | | | | | | | | | | | | + | | | | |
| U296C | I23I | | | | | + | | | | | | | | | | | | | | | | | | | | | | | | | | | |
| A305G | E26E | | | | | | | | | | | | | | | | | | | | | | | | + | | | | | | | | |
| U348C | Y41H | | | | | | | | | | | | | | | | | + | | | | | | | | | | | | | | | |
| A429G | I68V | | | | | | | | | | | | | | + | + | | | | | | | | | | | | | | | | | |
| A450G | I75V | | | | | | | | | | | | | | | | | | | | | | | | | + | | + | | | | | |
| G468U | D81Y | | | | | | | | | | | | | + | | | | | | | | | | | | | | | | | | | |
| A483G | I86V | | | | | | | | | | | | | | + | | | | | | | | | | | | | | | | | | |
| U504A | W93R | | | | | + | | | | | | | | | | | | | | | | | | | | | | | | | | | |
| C520U | A98V | | | | | | | | | | | | | | | | | | | | | | | | | | | | + | | | | |
| A524G | A99A | | | | + | | | | | | | | | | | | | | | | | | | | | | | | | | | | |
| A546G | N107D | | | | | | | | | | | | | | | | | | | + | | | + | | | | | | + | | | | |
| A561G | R112G | | | | + | | | | | | | | | | | | | | | | | | | | | | | | | | | | |
| U566C | P113P | | | + | | | | | | | | | | | | | | | | | | | | | | | | | | | | | |
| U583C | F119S | | | | | | + | | | | | | | | | | | | | | | | | | | | | | | | | | |
| U620C | A131A | | | | | | | | | | | | + | | | | | | | | | | | | | | | | | | | | |
| -626A | INSERT | + | | | | | | | | | | | | | | | | | | | | | | | | + | | + | | | | | |
| A642C | I139L | | | | | + | | | | | | | | | | | | | | | | | | | | | | | | | | | |
| A749G | A174A | | | | | | + | | | | | | | | | | | | | | | | | | | | | | | | | | |
| C756U | Q177STOP | | | | | | | | | | | | | | | | | + | + | | | | | + | | | | | | + | | | |
| U777C | L184L | | | | | | | | | | | | | | | | | | | | | | | | | | | | | | + | | |
| U787- | DELETION | | | | | | | | | | | | | | | | | | | + | | | | | | | | | | | | | |
| C808U | T194I | | | | | | | | | | | | | | | | | | | | | + | | | | | | | | | | | + |
| A825G | I200V | | + | | | + | + | | | | | + | | + | | | | + | + | + | + | + | + | + | + | + | + | + | + | + | + | + | + |
| A867G | K214E | | | | | | | | | | + | | | | | | | | | | | | | | | | | | | | | | |
| A873G | S216G | | | | | | | | | | | | | | | | + | | | | | | | | | | | | | | | | |
| U918C | F231L | | | | | | | | | | | | | | | | | | | | | | | | | | | | + | | | | |
| U963C | C246R | + | + | | + | + | + | + | + | + | + | + | | + | + | | + | + | + | + | + | + | + | + | + | + | + | + | + | + | + | + | + |
| G964A | C246H | | + | | | | | | | | | | | | | | | | | | | | | | | | | | | | | | |
| U1025G | V266V | | | + | | | | | | | | | + | | | + | | | | | | | | | | | | | | | | | |
| A1040G | A271A | | | | | | | | | | | | | | | | | | | | | + | | | | | | | | | | | + |
| A1061G | A278A | | | | | | | | | | | | | | | | | | | | | | | | | | | | | | | + | |
| U1070C | S281S | | | | | | | | | | | | | | | + | | | | | | | | | | | | | | | | | |
| A1106G | R292R | | | | | | | | | | | | | | | | + | | | | | | | | | | | | | | | | |
| C1130U | D301D | | | | | + | | | | | | | | | | | | | | | | | | | | | | | | | | | |
| U1153C | L309S | | | | | | | | + | | | | | | | | | | | | | | | | | | | | | | | | |
| A1211G | T328T | | | | | | | | | | | | | | | | | | | + | | | + | | | | | | + | | | | |
| U1231- | DELETION | | | + | | | | | | | | | | | | | | | | | | | | | | | | | | | | | |
| U1295C | V356V | | | + | | | | | | | | | + | | | | | | | | | | | | | | | | | | | | |
| U1349C | V374V | + | | | + | | | + | + | + | + | | | + | + | | + | | | | | | | | | | | | | | | | |
| A1353U | K376STOP | | + | | | | | | | | | | | | | | | | | | | | | | | | | | | | | | |
| U1391C | F388F | | + | | | | | | | | | | | | | | | | | | | | | | | | | | | | | | |
| U1392A | S389T | | | + | | | | | | | | | + | | | | | | | | | | | | | | | | | | | | |
| G1419A | G398S | | | | | | | | | | | | | | | | + | | | | | | | | | | | | | | | | |
| A1432G | Y402C | | + | | | | | | | | | | | | | | | | | | | | | | | | | | | | | | |
| C1483U | P419L | | | | | | | | | | | | | | | | | | | | + | | | | | | + | | | | | | |
| U1501G | V425G | | | | | | | | | | | | | | | | | | | | | | | | + | | | | | | | | |
| U1513C | L429P | | | | | | | | | | | | | | | | | | | | | | | | | | | | | | | + | |
| G1519A | R431Q | | | | | | | | | | | | | | | | | | | | | + | | | | | | | | | | | |
| A1534G | D436G | | | | | | | | | | | | | | | | | + | + | | | | | + | | | | | | + | | | |
| G1554A | A443T | | | | | | | + | | | | | | | | | | | | | | | | | | | | | | | | | |
| U1562C | S445S | | | | | | | | + | | | | | | | | | | | | | | | | | | | | | | | | |
| A1604G | Q459Q | + | | | + | | | + | + | + | + | | | + | + | + | + | | | | | | | | | | | | | | | | |
| G1635- | DELETION | | | | | | | | | | | | | | | | | | | | | | | | | | | | | | + | | |

**S2 Table. The list of mutations in the Rep- and RepNDK-RNA clones obtained at round 79 of the long-term replication experiment shown in Fig 3A. (Continued)**

| | | Rep-RNA clones | | | | | | | | | | | | | | | | RepNDK-RNA clones | | | | | | | | | | | | | | | |
|---|---|---|---|---|---|---|---|---|---|---|---|---|---|---|---|---|---|---|---|---|---|---|---|---|---|---|---|---|---|---|---|---|---|
| | | 1 | 2 | 3 | 4 | 5 | 6 | 7 | 8 | 9 | 10 | 11 | 12 | 13 | 14 | 15 | 16 | 1 | 2 | 3 | 4 | 5 | 6 | 7 | 8 | 9 | 10 | 11 | 12 | 13 | 14 | 15 | 16 |
| G1638A | A471T | | | | | | | | | + | | | | | | | | | | | | | | | | | | | | | | | |
| -1674A | INSERT | | | | | | | | | | | | | | | | | | | | | | | | | | | | | | | | + |
| G1698A | V491I | | | | + | | | | | | | | | | | | | | | | | | | | | | | | | | | | |
| G1725A | D500N | | | | + | | | + | | | | | | | | | | | | | | | | | | | | | | | | | |
| A1729G | Q501R | | | | | | | | | | | | | | | | | | | | | | | | | | | | | | + | | |
| A1759G | Y511C | | + | | | | | | | | | | | | | | | | | | | | | | | | | | | | | | |
| A1762G | D512G | + | | | | | | | | | | | | | | | | | | | | | | | | | | | | | | | |
| U1776C | C517R | | + | | | + | + | | | | | + | | | | | | + | + | + | + | + | + | + | + | + | + | + | + | + | + | + | + |
| U1801A | L525STOP | | | | | | | | | | | | | | | | | | | | | | | | | + | | + | | | | | |
| U1814C | G529G | | | | | | | | | | | | | | | | | + | + | | | | | + | | | | | | + | | | |
| U1972C | | | | | | | | | | | | | | | | | | + | + | | | | | + | | | | | | + | | | |
| C2000U | | | | | | | | | | | | | | | | + | | | | | | | | | | | | | | | | | |
| C2028U | | Not analyzed (Linkage region) | | | | | | | | | | | | | | | | | | | | + | | | | | | | | | | | + |
| G2030C | | Not analyzed (Linkage region) | | | | | | | | | | | | | | | | | | | | + | | | | | | | | | | | + |
| -2030G | | Not analyzed (Linkage region) | | | | | | | | | | | | | | | | | | | | | | | | | + | | | | | | |
| G2036- | | Not analyzed (Linkage region) | | | | | | | | | | | | | | | | | | | | | | | | | | | | + | | | |
| U2037G | | Not analyzed (Linkage region) | | | | | | | | | | | | | | | | + | | | | + | | | | | | | | | | | + |
| U2037- | | Not analyzed (Linkage region) | | | | | | | | | | | | | | | | | | | | | | | | | | | | | | | |
| C2038U | | Not analyzed (Linkage region) | | | | | | | | | | | | | | | | + | | | | + | | | | | | | | | | | + |
| C2041- | | Not analyzed (Linkage region) | | | | | | | | | | | | | | | | | + | | | | | + | | | | | | + | | | |
| -2042C | | Not analyzed (Linkage region) | | | | | | | | | | | | | | | | | | + | | | + | | + | | | | + | | + | + | |
| -2042CCC | | Not analyzed (Linkage region) | | | | | | | | | | | | | | | | | | | | | | | | | | | | | + | | |

**S3 Table. The list of mutations in the NDK- and RepNDK-RNA clones obtained at round 79 of the long-term replication experiment shown in Fig 3A.**

| | | NDK-RNA clones | | | | | | | | | | | | | | | | RepNDK-RNA clones | | | | | | | | | | | | | | | |
|---|---|---|---|---|---|---|---|---|---|---|---|---|---|---|---|---|---|---|---|---|---|---|---|---|---|---|---|---|---|---|---|---|---|
| | | 1 | 2 | 3 | 4 | 5 | 6 | 7 | 8 | 9 | 10 | 11 | 12 | 13 | 14 | 15 | 16 | 1 | 2 | 3 | 4 | 5 | 6 | 7 | 8 | 9 | 10 | 11 | 12 | 13 | 14 | 15 | 16 |
| G1A | | Not analyzed (Linkage region) | | | | | | | | | | | | | | | | | | | | + | | | | | | | | | | | + |
| G3A | | | | | | | | | | | | | | | | | | + | + | + | + | | + | + | + | | + | | + | + | + | + | |
| C6A | | | | | | | | | | | | | | | | | | | | | | + | | | | | | | | | | | + |
| U12G | | | | | | | | | | | | | | | | | | + | + | + | + | + | + | + | + | | + | | + | + | + | + | + |
| G15A | | | | | | | | | | | | | | | | | | | | | | | | | | | | + | | | | | |
| A133- | | | | | | | | | | | | | | | | | | | | | + | | | | | | + | | | | | | |
| G138A | | | | | | | | | | | | | | | | | | | | | | | | | | + | | + | | | | | |
| A205G | | | | | | | | | | + | | | | | | | | | | | | | | | | | | | | | | | |
| C228U | | | | | | | | | | | | | | + | | | | + | + | | | + | | + | + | | | | | + | | | + |
| U237C | I3I | | | | | | | | + | | | | | | | | | | | | | | | | | | | | | | | | |
| A240G | E4E | + | + | | + | | | | | + | | + | | | + | + | | | | | | | | | | + | | + | | | | | |
| A244G | T6A | | | + | | | | | | | | | | | | | | | | | | | | | | | | | | | | | |
| U246C | T6T | | | | | | | | | | | | | | | | | | | | | | | | | | | | | | | | + |
| U324G | F32L | | | | | | | | | | | | | | | | | | | | | | | | + | | | | | | | | |
| U333C | V35V | + | | | | | | | | | | | | | | | | | | | | | | | | | | | | | | | |
| U414C | G62G | | | + | | | | | | | | | | | | | | | | | | | | | | | | | | | | | |
| U465C | V79V | + | + | | + | | | | | + | | + | | | + | + | | + | + | | | | | + | | + | | + | | + | | | |
| G475A | V83I | | | | | | | | | | | | | | | + | | | | | | | | | | | | | | | | | |
| U477C | V83V | | | | | + | | | | | | | | | | | | | | | | + | | | | | | | | | | | |
| G498A | L90L | + | + | | + | | | | | + | | + | | | + | + | | | | | | | | | | + | | + | | | | | |
| U582C | G118G | + | + | | + | | + | + | + | + | | + | + | | + | + | | | | | | | | | | + | | + | | | | | |
| A603G | A125A | | | | | | | | | | + | | | | | | | | | | | | | | | | | | | | | | |
| U623C | F132S | | | | | | | | | | | | + | | | | | | | | | | | | | | | | | | | | |

**S4 Table. The list of primers (from 5′ end to 3′end).**

| | |
|---|---|
| Primer1 | Rep_F: ATACACATGGCTCGTAGAAAA |
| Primer2 | Rep_R: GGCGTACACGCTTGCGGAAGT |
| Primer3 | NDK_F: CCATCATTAAACCAAATGCAGTAGC |
| Primer4 | NDK_R: GGTTTTCCATCGTGTTCAGC |
| Primer5 | RepNDK_F: GTGCTGTTGTTCGGGTTGTTG |
| Primer6 | RepNDK_R: GATGGAAAAAGTACGTTCAATCGCC |
| Primer7 | Rep_or_NDK_F: GGGTCACCTCGCGCAGC |
| Primer8 | Rep_or_NDK_R: CCGGAAGGGGGGGACGAGG |
| Primer9 | RepNDK_F2: AAGACAGCATCTTCGCGTAACTCTC |
| Primer10 | RepNDK_R2: CGGGCACACCTCACCTTCTCCAAAG |
| Primer11 | Vector_F: GTCCCCCCCTTCCGGGGGGGTCCCC |
| Primer12 | Vector_R: TGCGCGAGGTGACCC |
| Primer13 | NDKRep_F: ATATACACATGGCGATTGAACGTAC |
| Primer14 | NDKRep_R: CAAGCCTAACAACAACCCGAAC |
